# Environmental and anthropic factors influencing *Aedes aegypti* and *Aedes albopictus* (Diptera: Culicidae), with emphasis on natural infection and dissemination: Implications for an emerging vector in Colombia

**DOI:** 10.1101/2024.10.07.616949

**Authors:** Mantilla-Granados Juan S, Montilla-López Karol, Sarmiento-Senior Diana, Chapal-Arcos Elver, Velandia-Romero Myriam Lucía, Calvo Eliana, Morales Carlos Andrés, Castellanos Jaime E

## Abstract

**Background:** Viruses such as dengue virus (DENV), Zika virus (ZIKV), and chikungunya virus (CHIKV) pose major threats to human health, causing endemic, emerging, and reemerging diseases. These arboviruses have complex life cycles involving *Aedes* mosquitoes, driven by environmental, ecological, socioeconomic, and cultural factors. In Colombia, *Aedes aegypti* is the primary vector, but *Aedes albopictus* is expanding across the country. Understanding the unique characteristics of each species is crucial for managing arbovirus spread, particularly in areas where they coexist.

**Methodology/Principal Findings:** We conducted an entomological survey of *A. aegypti* and *A. albopictus* (larvae, pupae, and adults) in urban and rural areas of four municipalities across different elevations (200–2200 meters above sea level) in Colombia. Household conditions and knowledge of DENV were assessed through interviews. Female *A. albopictus* were tested individually for arbovirus RNA, while *A. aegypti* were tested in pools. Both species were found up to 2100 masl. *A. aegypti* comprised 78% of the immature forms collected, while *A. albopictus* made up 22%. Larvae from both species coexisted in common artificial breeding sites in urban and rural areas, with no evidence of competition. *A. albopictus* preferred rural areas, lower elevations (<1500 masl), high precipitation (>270 mm), and poor household conditions, while *A. aegypti* was more abundant in urban areas, intradomicile environments, and areas with moderate precipitation (100–400 mm). Biting risk was higher for *A. aegypti* (0.02–0.22 females per person), particularly in urban areas, while *A. albopictus* exhibited lower biting risk (0.001–0.08), with the highest values in rural Patía. Natural infections of DENV (12.4%) and CHIKV (12.4%) were found in *A. aegypti*, while *A. albopictus* showed CHIKV (41.2%) and DENV (23%) infections, with virus dissemination to the legs and salivary glands.

**Conclusions/Significance:** Integrating household conditions and community knowledge with environmental data can enhance predictive models for vector presence and guide surveillance and educational strategies. Our findings highlight the need to consider *A. albopictus* as a potentially significant arbovirus vector in Colombia, especially given the presence of arboviruses in its salivary glands, its use of artificial breeding sites, its biting risk inside homes, and its differing ecological preferences and seasonal associations compared to *A. aegypti*.

**AUTHOR SUMMARY:** In this study, we investigated the ecological and epidemiological dynamics of *Aedes aegypti* and *Aedes albopictus* and their natural infection with DENV, ZIKV, and CHIKV in Colombia. An entomological survey conducted across four municipalities revealed distinct environmental and human-related factors influencing the distribution and abundance of these species. *A. aegypti* was more abundant in urban environments, favoring areas with lowest precipitation and a range of household conditions, while *A. albopictus* was more common in rural areas with higher precipitation and poorer household conditions. Both species were naturally infected with DENV and CHIKV, with *A. albopictus* showing the ability to disseminate, as indicated by their presence in the legs and salivary glands. Our findings underscore the importance of understanding species-specific ecological characteristics and incorporating community knowledge into predictive models to improve vector surveillance and control strategies, particularly in regions where both species coexist and contribute to arbovirus transmission.

## INTRODUCTION

Arboviruses account for > 17% infectious diseases worldwide, with a high proportion of emerging and zoonotic pathogens concentrated in tropical and subtropical regions [1]. Dengue virus (DENV) is the most significant arboviral threat in the Americas, causing the highest morbidity and mortality, and is endemic across tropics and subtropics. In 2024, the Pan American Health Organization has reported a 235% increase in DENV cases compared to that in 2023, and a 431% increase over the 5-year average [2]. The introduction of chikungunya virus (CHIKV) and Zika virus (ZIKV) to the Americas in 2013 and 2015, respectively, has also raised major public health concerns owing to their association with chronic diseases, neurological effects, and congenital infections [3–5]. Despite vector control efforts, the incidence of these other arboviruses is increasing in the region, likely driven by climate change and unplanned urbanization [6]. Shifts in incidence may also result from changes in the epidemiological landscape owing to the introduction of new competent vectors.

The primary vector for these arboviruses is the African mosquito, *Aedes aegypti* (L.), which has long been established in the Americas. It breeds in artificial containers, mainly in urban areas; however, it can also colonize rural settings [7]. The Asian tiger mosquito *Aedes albopictus* is a highly adaptable species, which can use both natural and artificial larval habitats, allowing it to thrive in the tropical and temperate areas, and at altitudes higher than that for *A. aegypti* [8–10]. *A. albopictus* (Skuse), is a secondary or even primary vector for DENV, ZIKV, and particularly CHIKV, depending on the genetic background that influences its vector competence and interactions with the local microbiota [11,12].

Field-collected mosquitoes can be screened for the presence of arbovirus in the whole body, abdomen, and midgut to assess natural infection, which may reflect an ingested or disseminated virus. Virus detection in the legs or organs outside the midgut indicates dissemination, whereas infection in the salivary glands or saliva is associated with transmission potential [11,13].

*A. aegypti* and *A. albopictus* occupy similar niches and may compete, with *A. albopictus* sometimes displacing *A. aegypti* [14–16]. As invasive species are well adapted to human-modified environments, their presence can be shaped by climate and social factors, which could sustain arboviral transmission for long periods [17]. While both vectors are found in the Americas, *A. albopictus* is considered secondary because of its restricted distribution, relatively low population densities, and reduced vector competence in some populations [18,19]. However, many *A. albopictus* populations across Latin America remain understudied, and filling this knowledge gap is critical for designing improved surveillance and control strategies.

In Colombia, *A. albopictus* was first reported in 1998, and has since spread to 32 departments of the country. This mosquito is naturally infected by DENV, ZIKV, and CHIKV [20–25]whit some studies demonstrating its ability to colonize major urban areas [26]. However, the role of this species as a vector in Colombia, its relationship with *A. aegypti*, and the climatic and anthropogenic factors associated with their distribution are poorly understood.

The Cauca Department is an ideal study area for these issues, as both *Aedes* species have been widely reported there. The region experiences a high incidence of dengue, and various dengue control efforts through educational, physical, and chemical approaches have been implemented. Therefore, we aimed to investigate the presence of *A. aegypti* and *A. albopictus* concerning climatic and anthropogenic determinants in this setting, and their status of natural infection.

Overall, our results revealed distinct ecological patterns for *A. aegypti* and *A. albopictus* populations across different environmental and anthropic conditions in the surveyed Colombian municipalities. Arbovirus infections, including DENV and CHIKV, were detected in both species, with notable virus dissemination and salivary glands infections in *A. albopictus*, suggesting its role as arbovirus vector.

## METHODS

### Study area

This study was performed in the urban and rural areas of four municipalities of the Cauca Department, selected for sampling at certain altitudinal ranges: Piamonte [0–600 m above sea level (masl)], Patía (600–1200 masl), Piendamó (1200–1800 masl), and Popayán (1800–2200 masl) (Fig 1).

**Fig 1.**
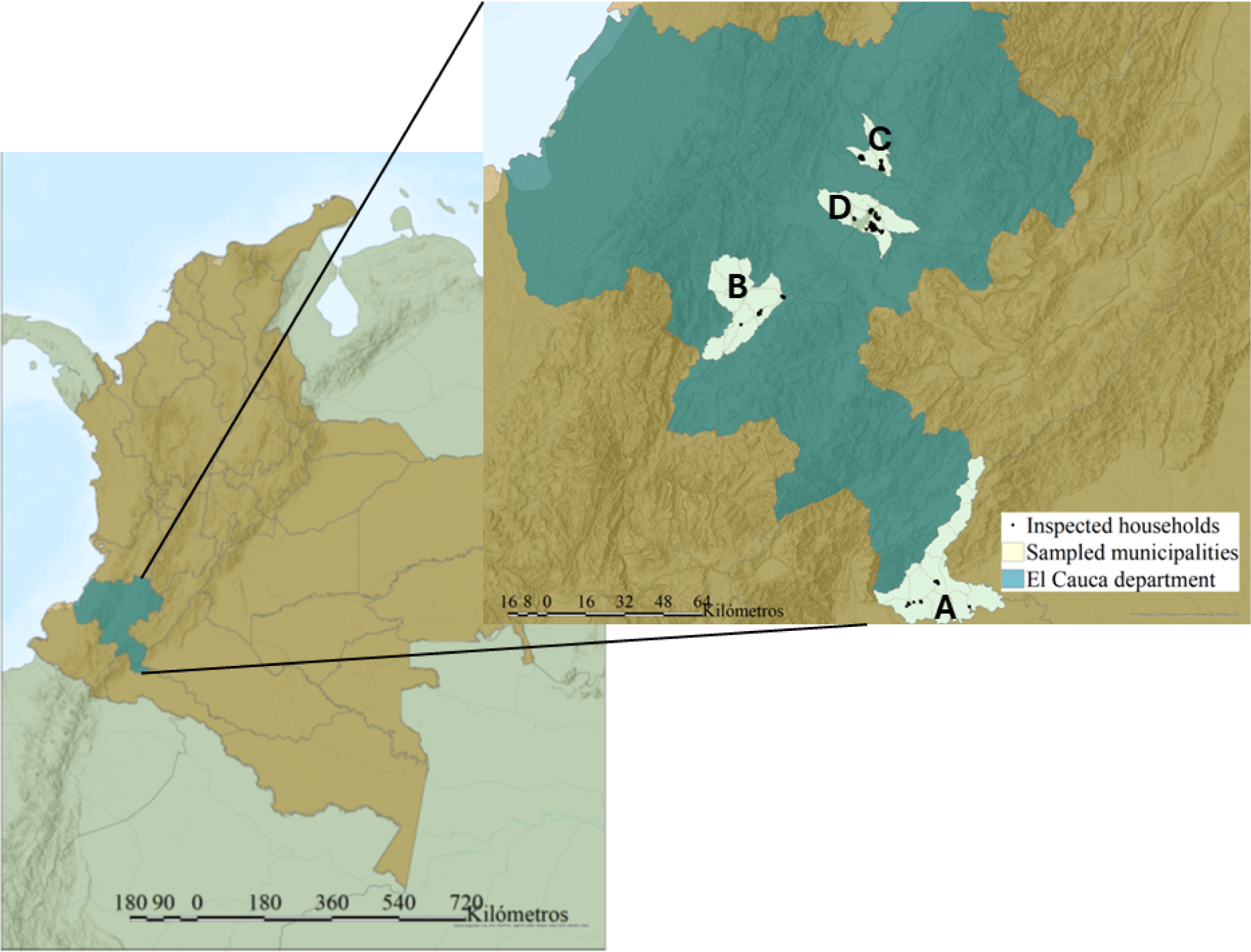
Map for the sampled localities in Colombian Map showing the localization of Cauca department in dark aquamarine. A–D Municipality locations A. Piamonte. B. Patía. C. Piendamó. D. Popayán. Black dots indicate the localization of sampled households.

### Sample size

This cross-sectional study used the household as the unit of analysis, with the sample size determined using projected population data from the national census (https://www.dane.gov.co/) and calculated using SSPropor tool (http://www.openepi.com/), each house was visited twice for mosquitoes collection during a dry (July-October 2021) and a wet (March-May 2022) periods.

### Approvals and ethical considerations

Before data collection, written informed consent was obtained from the participants. This study was approved by the Institutional Ethics Committee for Research at Universidad El Bosque, Bogotá, Colombia (act number: 013-2021). Mosquito sampling was authorized by the National Authority of Environmental Licenses (resolution number: 01470, November 17, 2017).

### Mosquito collection

Mosquito sampling was conducted between July–October 2021 and March–May 2022. Immature mosquitoes were collected directly from water containers of urban and rural settlements, which were categorized into artificial containers, including laundry sinks, ground tanks (plastic storage tanks with > 2000 L of capacity, kept on the floor), tires, buckets, drain sewers; others, including animal water bowls, flowerpots, cisterns, bottles, discarded plastic waste, cups, pots, plastics, caps, and tubes; and natural containers, including bamboos, aquatic plants, tree holes, and plant axils, with most of the categorization proposed in intradomicile, peridomicile (0–10 m from the household), and extradomicile (> 10 m from the household) areas [27]. Sunlight exposure of the breeding places was also recorded as uncover (direct sunlight), partial or cover (completely under the shadow). Pupae were allowed to emerge to adults. Adult resting mosquitoes were captured using a Prokopack aspirator [28] for 15 min each from intradomicile and peridomicile areas, and using entomological nets from extradomicile areas when they were observed. The mosquito collection sites were georeferenced using Garmin GPS 62s (Garmin International Inc., Olathe, KS, USA).

### Mosquito identification

Immature and adult mosquitoes were taxonomically identified in the field following the characteristics previously described [29,30] for each species, using a Zeiss ™ Stemi ™ DV4 (Jena, Germany). The identified *A. aegypti* specimens were pooled by life stage, sex, date, and location, and preserved in RNAlater™ Stabilization Solution (Ambion, AM7020) (Thermo Fisher Scientific, Waltham, Massachusetts, U.S.A.). Female *A. albopictus* was dissected into the midgut, legs, and thorax, and preserved individually in RNAlater. The males and immature mosquitoes of both species were processed similarly to that for *A. aegypti*.

### Entomological index

We calculated the following entomological indicators: percentage of key positive, the number of positive water containers of each category over the total number of positive containers for each species; Breteau index, the number of positive water-holding containers per 100 houses over the total inspected houses; pupae per person index, to assess the rate of contact between humans and potential infected vectors [31]; and females per person index, the number of females recollected on each house by one person during 15 min with a prokopack aspirator. For adult female mosquitoes, the infestation was estimated as the number of females collected over 15 min using one Prokopack trap within intradomicile areas. The total numbers of collected larvae and pupae of each species per house were also analyzed.

### Environmental variables

The average, minimum, and maximum temperature values and total precipitation were obtained from WorldClim data [32]. The geographic coordinates and elevations recorded in the field were projected and associated with environmental variables using ArcGIS Pro v.3.3.

### Anthropogenic variables

The residents of each sampled household were interviewed about various sociodemographic and economic characteristics, including access to running water (and its continuity), water storage and rainwater harvesting practices, trash disposal services and methods (burying or burning), and primary construction materials [concrete, wood, or other (like shade mesh walls)]. Additionally, general knowledge about dengue transmission, the most common arbovirus in the study area, was assessed, such as whether they had previously received information about dengue, understood the transmission mechanism, recognized who was at risk of DENV transmission, and could identify correct and incorrect *Aedes* breeding sites. These variables were recorded as binary (0/1) indicators of the presence or absence and grouped into two principal components using FactoMineR library in R v.4.3.3.

Among the previously considered variables, we retained those best represented in the first two components and increased the percentage of explanation of the data variance. The first component was dominated by household characteristics and labeled ‘house conditions’, with some of the variables as previously described in [33] as ‘environmental capital’; we retained the availability of running water, sewerage, and the houses made of concrete as the main variables associated with houses with high values on this component (house conditions), while the ones with negative values were without these public services, use of wood as construction material and the practice of burning wastes, houses scored as cero, and medium house conditions (availability of some of the public services). The second component was characterized by knowledge variables on dengue and vector and labeled ‘knowledge’ (S1 Fig); the variables retained for this component were reports to have received previous information about dengue and its vector, correct recognition of potential breeding places for dengue vectors, and association of dengue transmission with mosquito bite, and positive values for the houses were considered with correct recognition of these categories; Negative scores were associated with incorrect information on these categories, and cero values corresponded to houses with inhabitant unaware about these topics. The other variables were not used for the final principal component analysis because of their low representation of the first two components, decreasing their explanatory percentage on the data variance; however, the general pattern of the total variables examined at the beginning was maintained with the selected variables (S1 Fig). The first two principal components explained 30.27% and 22.7% of the data variance. The coordinates of each household along these principal components were used as response variables for statistical analyses.

### Statistical analysis

To analyses larvae abundance of each species, related to breeding places characteristics and abundance of the other species. The analysis was performed by using a generalized linear model (GLM) with poisson distribution using function ‘glm’ from MASS [34] package in R program version 4.3.3. For female infestation (number of females collected per household in 15-min Prokopack aspiration), the number of larvae of each species was also analyzed against environmental variables, housing conditions and knowledge were separately evaluated because of collinearity. The presence of multicollinearity was assessed using the generalized variance inflation factor GVIF^(1/(2×Df))^, with the ‘car’ package [35] in R 4.3.3 The data exhibited nonlinear patterns with the response variables; therefore, generalized additive models (GAMs) with negative binomial distributions were implemented using mgcv package[36] in R v.4.3.3 Cubic regression spline smoothing was applied with the k parameter manually adjusted for each response variable.

### RNA extraction and amplification

RNA was extracted from mosquito samples using a Viral Nucleic Acid Extraction Kit II (IBI Scientific, Chavenelle, Dubuque, U.S.A.), according to the manufacturer’s protocol, and quantified using a RNA samples were quantified using the Nanodrop system NanoPhotometer® NP80 (Implen, Westlake Village, California, U.S.A.). Viral RNA was reverse-transcribed and amplified by multiplex nested reverse transcriptase–polymerase chain reaction using a Luna® Universal kit One-Step RT-PCR (Thermo Fisher Scientific, Waltham, Massachusetts, U.S.A.) and primers for DENV, CHIKV, and ZIKV, as previously described [37]. Culture-harvested viruses were used as positive controls. Amplified products were separated on 2% agarose gels and visualized by staining with ethidium bromide. For *A. albopictus* samples, abdominal or midgut amplifications were recorded as natural infections, whereas leg/thorax or salivary gland amplifications indicated viral dissemination, the percentage of positive pools or individuals (complete or dissected) was calculate, in the case of pools the minimum infection rate (MIR) was calculated as the number of positive pools / total specimens tested x 1000.

## RESULTS

In total, 904 households were sampled across urban (n = 566) and rural (n = 338) areas (Table 1). These households were visited twice (during the dry and wet seasons); however, not all households were visited twice because of residential unavailability. The final sample sizes according to municipality and season are listed in Table 1.

**Table 1.**
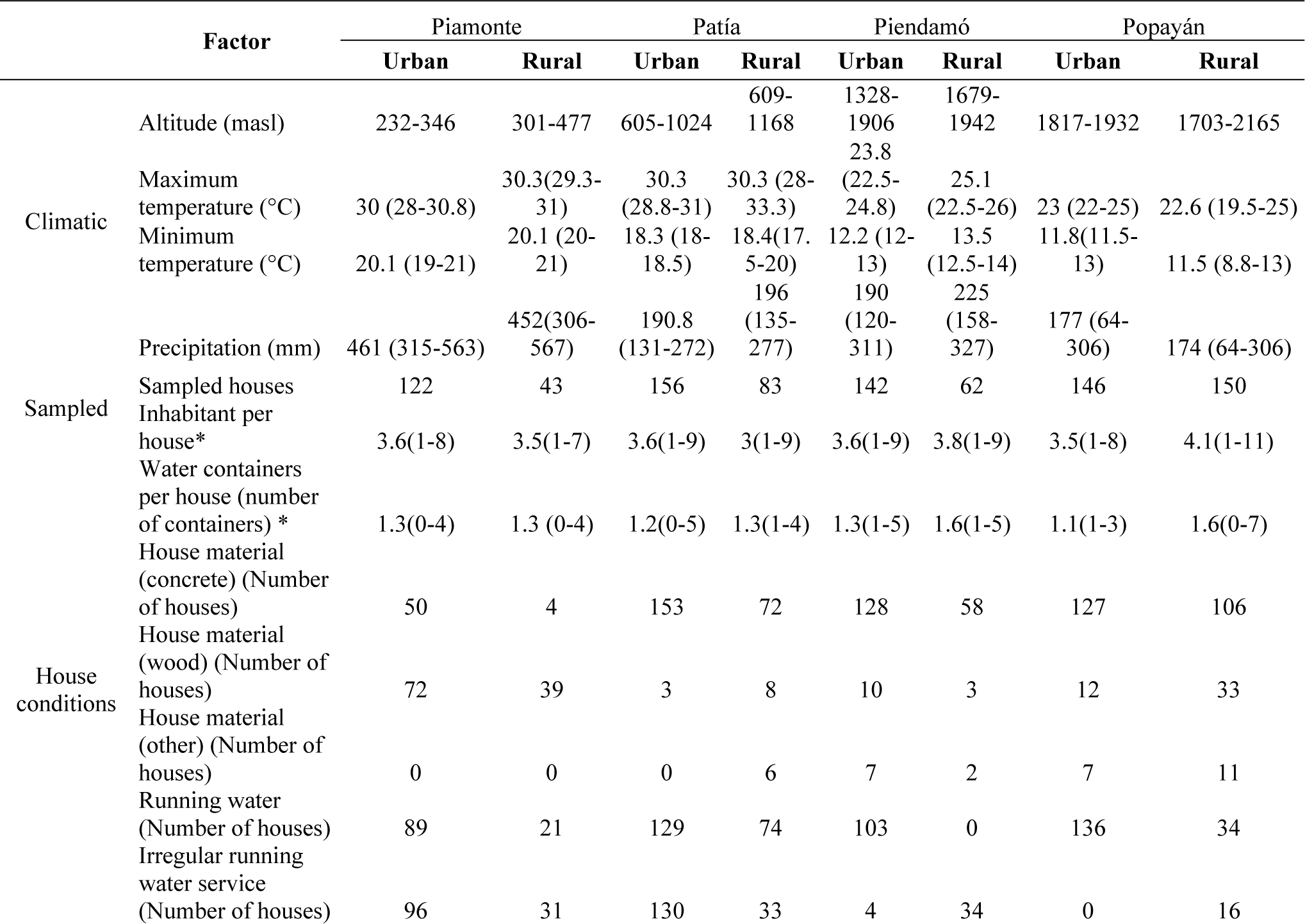
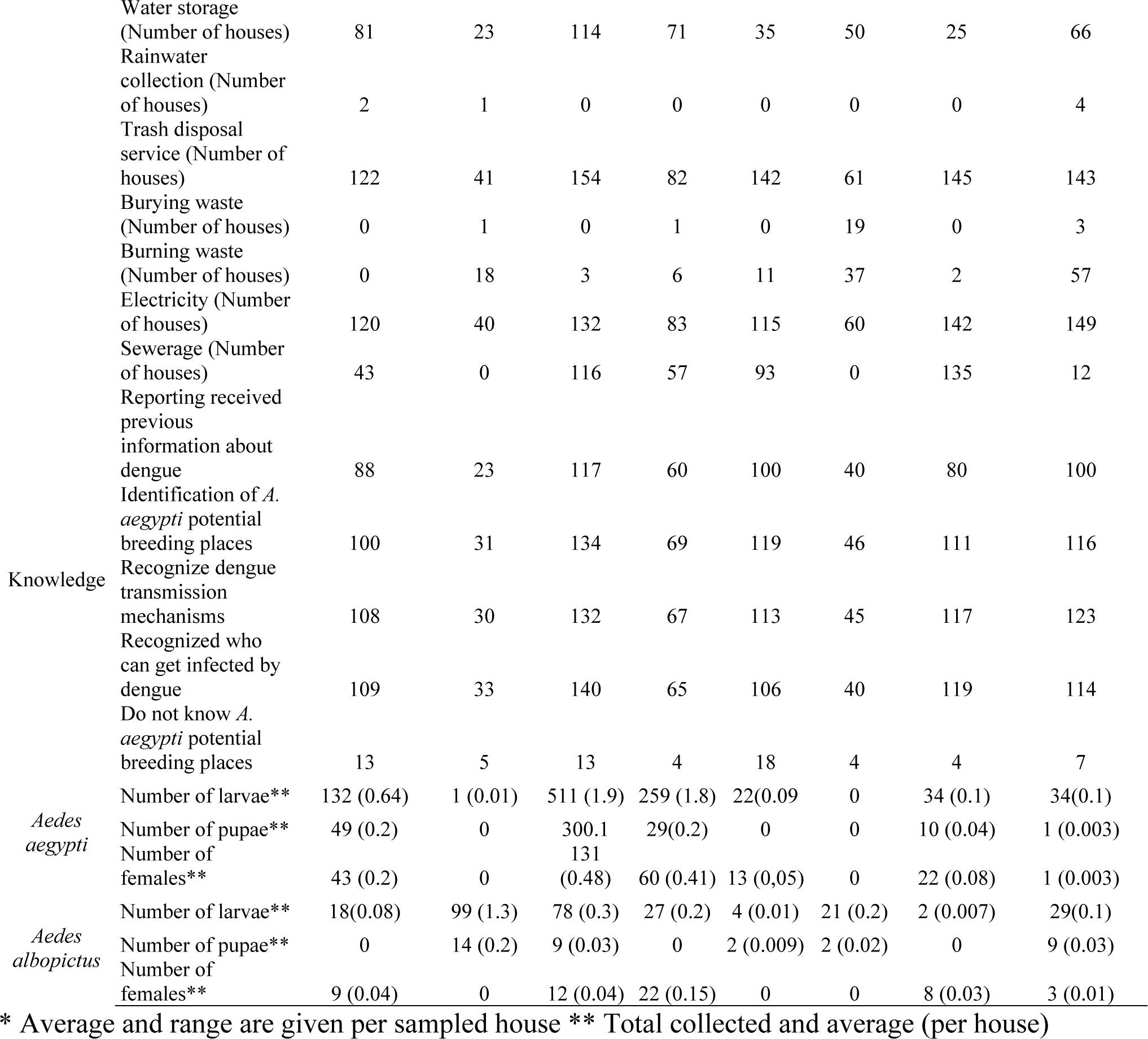
Summarized data of urban and rural areas per each sampled municipality.

Differences in several parameters were observed between urban and rural households: the main construction material used for urban houses was concrete 81% (458), followed by wood 17% (97), and other materials like shade mesh 2% (14); at the rural area, 71% (240) houses were made of concrete, 24% (83) of wood, and 5.6% (19) of plastic shade mesh. A total of 457 (80.7%) and 129 (38.1%) of the urban and rural houses, respectively, reported access to running water; however, 50% (230) and 88.4% (114) reported experiencing irregular services in urban and rural areas, respectively. The practice of water storage was reported in 45% (255) urban and 62.1% (210) rural houses. Waste disposal services were available to 563 (99.4%) urban and 327 (96.7%) rural houses; burying trash was reported in only 7.1% (24) rural houses; and burning was reported in 2.8% (16) urban and 34.9% (118) rural houses. Electricity access was reported in 89.9% (509) urban and 98.2% (332) rural houses, and sewerage in 68.4% (387) urban and 20.4% (69) rural houses.

Residents of 385 (68%) urban and 223 (66%) rural houses reported receiving prior information on dengue and its vector. Mosquito biting as the transmission mechanism was recognized by 83% (470) urban and 78.4% (265) rural residents. *A. aegypti* breeding sites were identified by 82% (464) urban and 78.4% (265) rural residents. Finally, 474 (83.7%) urban and 252 (74.5%) rural residents recognized that anyone could be infected with dengue.

### Immature stages

A total of 1,426 immature (larvae and pupae) *Aedes* mosquitoes were collected across the four municipalities, with 78% (1,112) *A. aegypti* (29% rural and 71% urban) and 22% (314) *A. albopictus* (64% rural, 35% urban) during the dry and wet seasons (Table 1). Both species were found in urban and rural areas, except in Piendamó where *A. aegypti* was detected only in the urban area. The Breteau index (BI), was highest for *A. aegypti* in Patía (BI = 18.2) followed by those in Piamonte (8.4), Popayán (1.6), and Piendamó (1.1). For *A. albopictus*, Patía had the highest BI (4.8), followed by those of Piamonte (2.5), Popayán (1.5), and Piendamó (0.9).

Despite this, we searched for natural and artificial containers in both intra and extra-domicile areas, such as parks, rivers, and closed forested areas, and found some natural water containers in both urban and rural areas (S1 Table). Both *Aedes* species primarily utilized artificial containers as larval habitats (Fig 2, S1 Table); however, specific container types varied between species and urban/rural settings. For *A. aegypti,* laundry sinks and buckets were the most common breeding sites in both urban and rural areas. In contrast, *A. albopictus* often used tires and laundry sinks in rural areas, whereas buckets and laundry sinks were preferred in urban settings. Notably, immature mosquitoes of both species were found in urban drain sewers, and *A. albopictus* used bamboo canes, although that were not the predominant breeding place (Fig 2).

**Figure 2.**
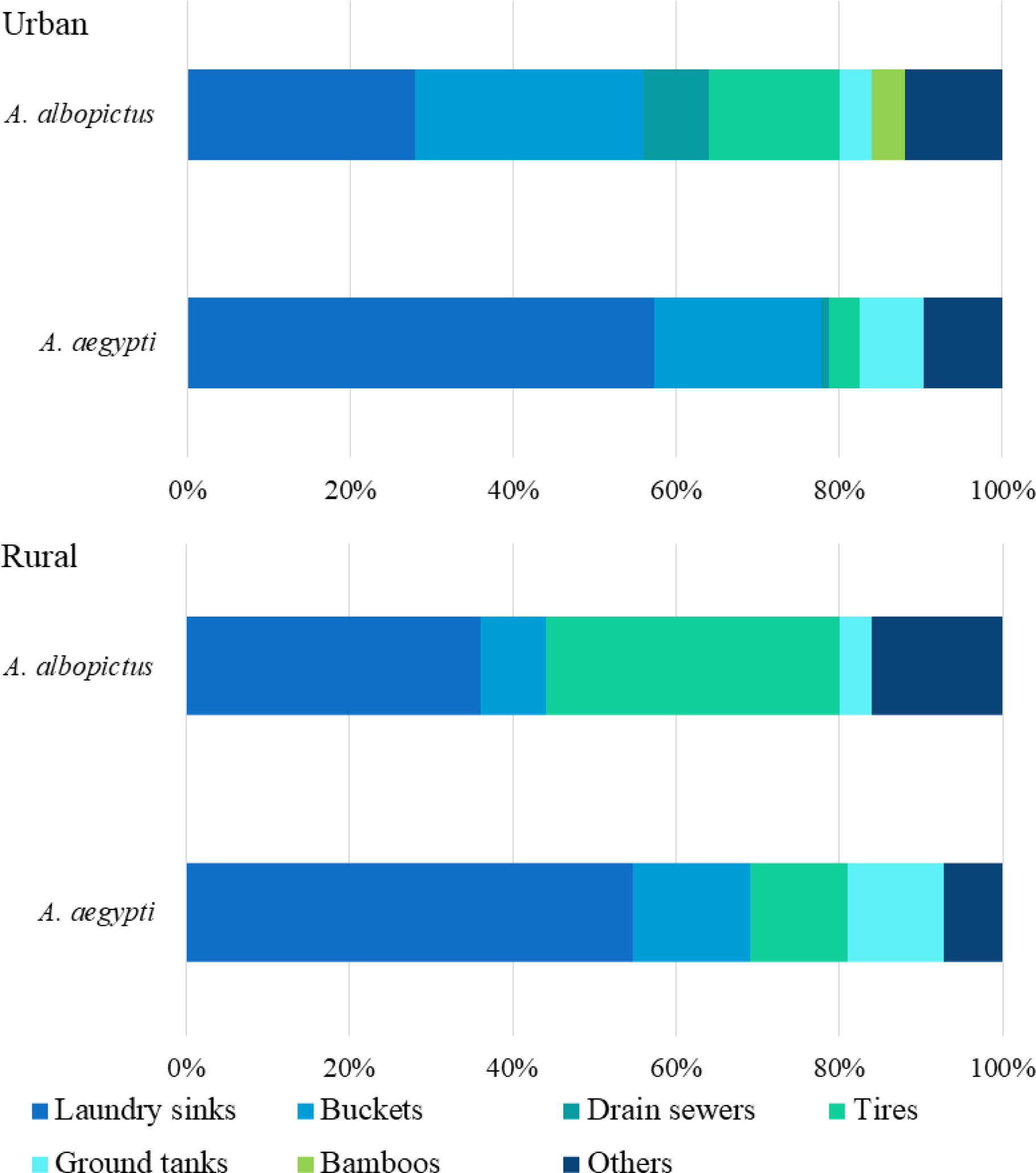
Percentage of key breeding places productivity used in urban and rural areas by *A. aegypti* and *A. albopictus*.

Both *A. aegypti* and *A. albopictus* were found up to 2,100 masl (Popayán rural settlement). Generalized linear models revealed that larval abundance of *A. aegypti* was significantly and positively associated with the number of collected *A. albopictus* larvae in the same container, urban areas, uncovered breeding places, and those located indoors. However, it was negatively associated with wet season and partially covered breeding places (Table 2). For *A. albopictus*, larval numbers were significantly linked with the number of *A. aegypti* larvae collected in the same container, wet season, partial cover, and breeding places located in the peridomicile area; however, they were negatively affected by urban areas and indoor breeding places (Table 2).

**Table 2.**
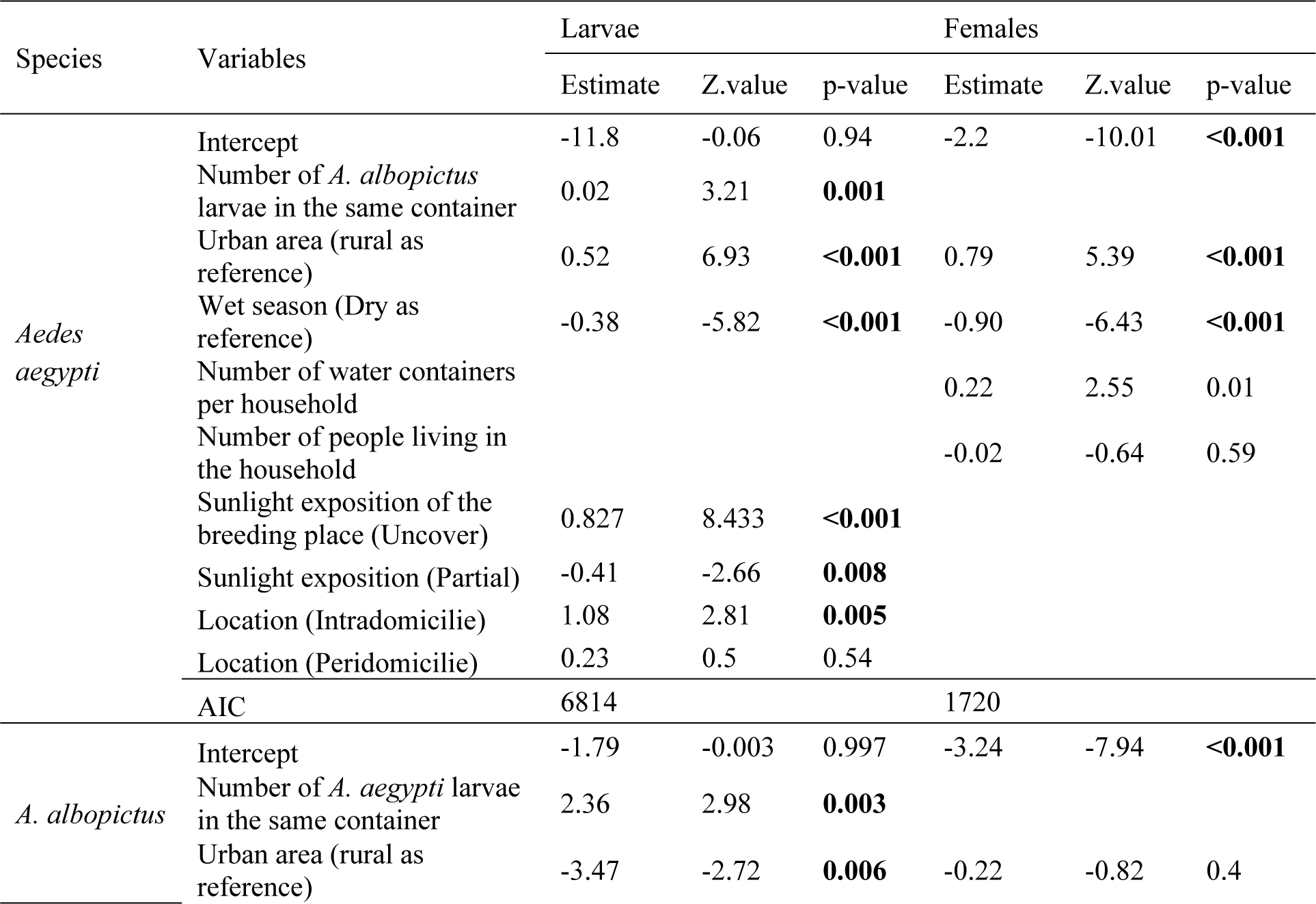
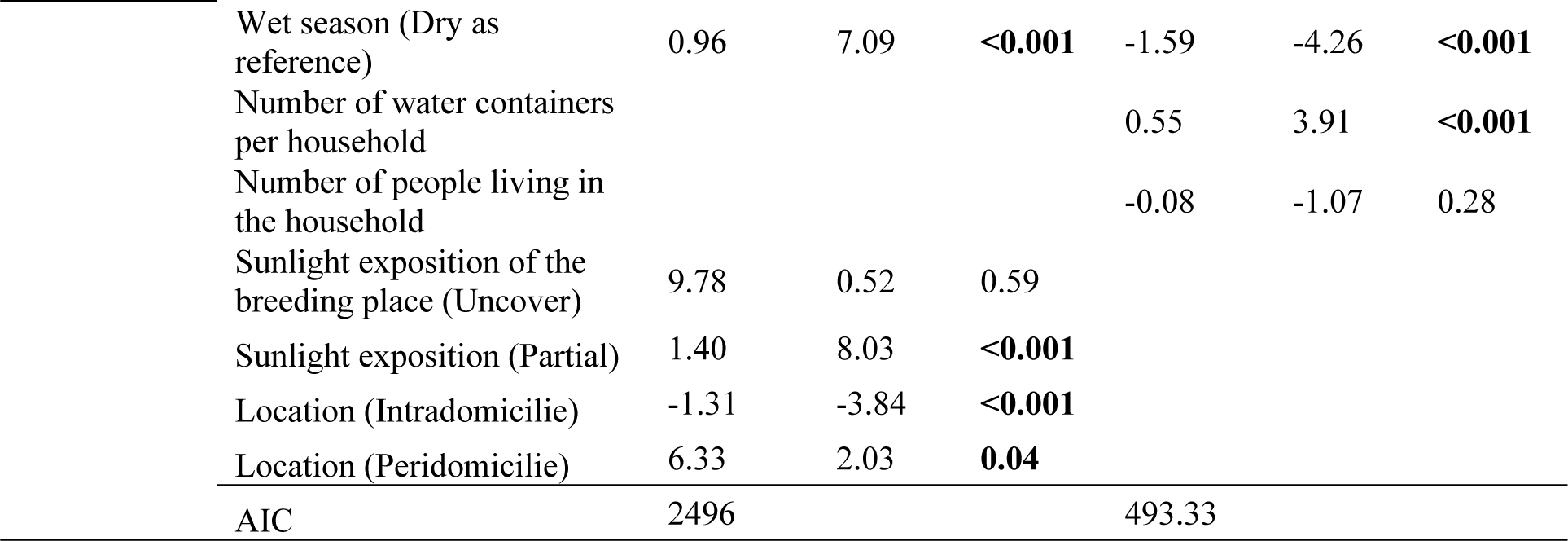
Analysis of the number of larvae and female mosquitoes regarding urban location, water containers, and inhabitants, by a generalized linear model (GLM) with Poisson distribution.

| Species | Variables | Larvae |  |  | Females |  |  |
| --- | --- | --- | --- | --- | --- | --- | --- |
|  |  | Estimate | Z.value | p-value | Estimate | Z.value | p-value |
| *Aedes aegypti* | Intercept | -11.8 | -0.06 | 0.94 | -2.2 | -10.01 | **<0.001** |
|  | Number of *A. albopictus* larvae in the same container | 0.02 | 3.21 | **0.001** |  |  |  |
|  | Urban area (rural as reference) | 0.52 | 6.93 | **<0.001** | 0.79 | 5.39 | **<0.001** |
|  | Wet season (Dry as reference) | -0.38 | -5.82 | **<0.001** | -0.90 | -6.43 | **<0.001** |
|  | Number of water containers per household |  |  |  | 0.22 | 2.55 | 0.01 |
|  | Number of people living in the household |  |  |  | -0.02 | -0.64 | 0.59 |
|  | Sunlight exposition of the breeding place (Uncover) | 0.827 | 8.433 | **<0.001** |  |  |  |
|  | Sunlight exposition (Partial) | -0.41 | -2.66 | **0.008** |  |  |  |
|  | Location (Intradomicilie) | 1.08 | 2.81 | **0.005** |  |  |  |
|  | Location (Peridomicilie) | 0.23 | 0.5 | 0.54 |  |  |  |
|  | AIC | 6814 |  |  | 1720 |  |  |
| *A. albopictus* | Intercept | -1.79 | -0.003 | 0.997 | -3.24 | -7.94 | **<0.001** |
|  | Number of *A. aegypti* larvae in the same container | 2.36 | 2.98 | **0.003** |  |  |  |
|  | Urban area (rural as reference) | -3.47 | -2.72 | **0.006** | -0.22 | -0.82 | 0.4 |

|  |  |  |  |  |  |  |
| --- | --- | --- | --- | --- | --- | --- |
| Wet season (Dry as reference) | 0.96 | 7.09 | **<0.001** | -1.59 | -4.26 | **<0.001** |
| Number of water containers per household |  |  |  | 0.55 | 3.91 | **<0.001** |
| Number of people living in the household |  |  |  | -0.08 | -1.07 | 0.28 |
| Sunlight exposition of the breeding place (Uncover) | 9.78 | 0.52 | 0.59 |  |  |  |
| Sunlight exposition (Partial) | 1.40 | 8.03 | **<0.001** |  |  |  |
| Location (Intradomicilie) | -1.31 | -3.84 | **<0.001** |  |  |  |
| Location (Peridomicilie) | 6.33 | 2.03 | **0.04** |  |  |  |
| AIC | 2496 |  |  | 493.33 |  |  |

The smooth curves from the GAM models indicated that larval numbers of *A. aegypti* increased at the smooth term for minimum temperature above 13 °C, peaked at 18 °C, and declined at 21 °C. Precipitation between 100–400 mm was also associated with high number of *A. aegypti* larvae, which reached the maximum value at approximately 250 mm and then decreased above 400 mm precipitation. Furthermore, larval numbers of *A. aegypti* increased with elevation from 100 to 1500 masl, peaking at 1000 masl (Fig 3A). In contrast, larval numbers of *A. albopictus* increased with minimum temperatures above 15 °C, and showed a strong positive relationship with the smooth term for precipitation above 270 mm, while elevations > 1500 masl reduced larval abundance (Fig 3A).

**Figure 3.**
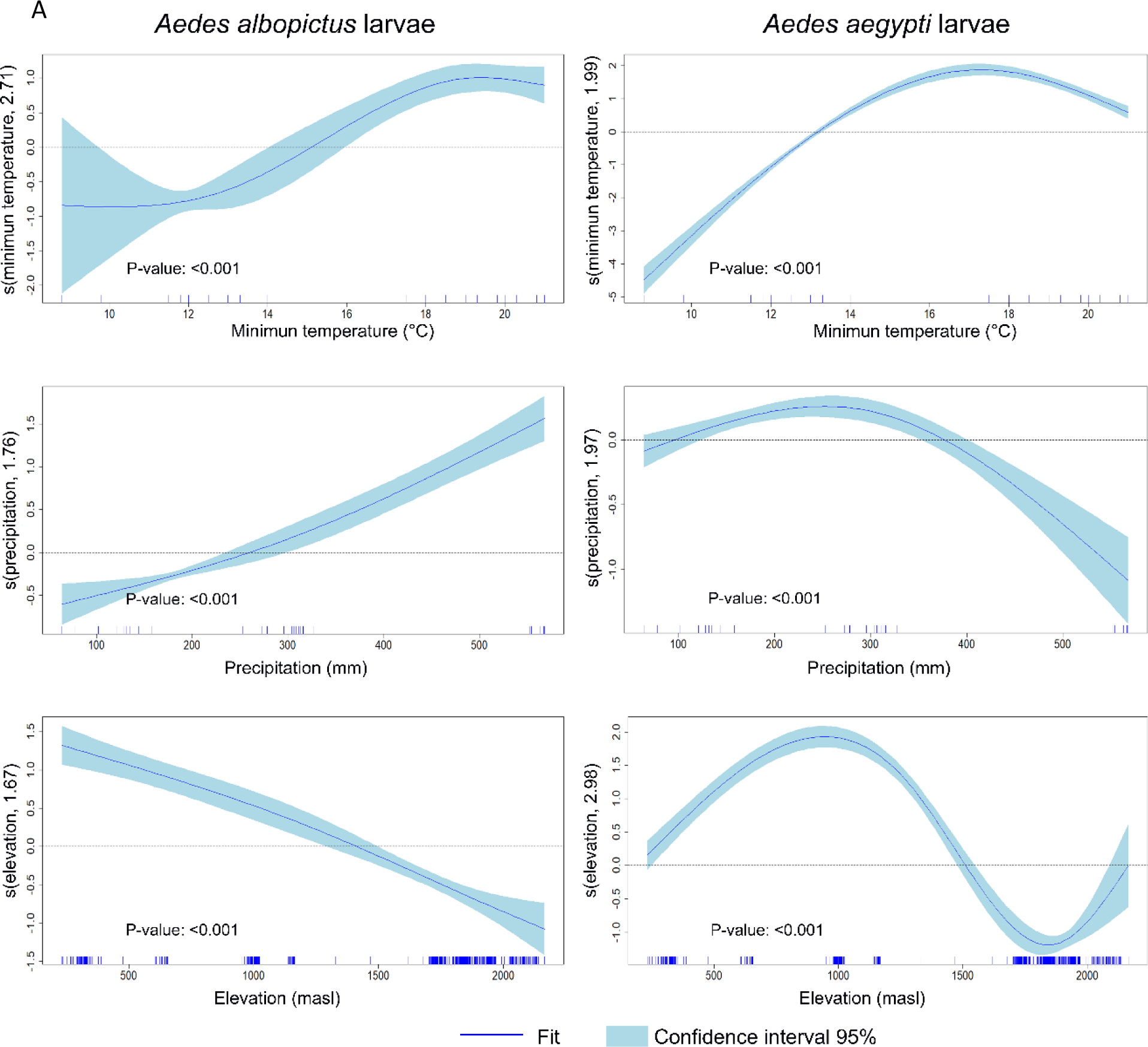
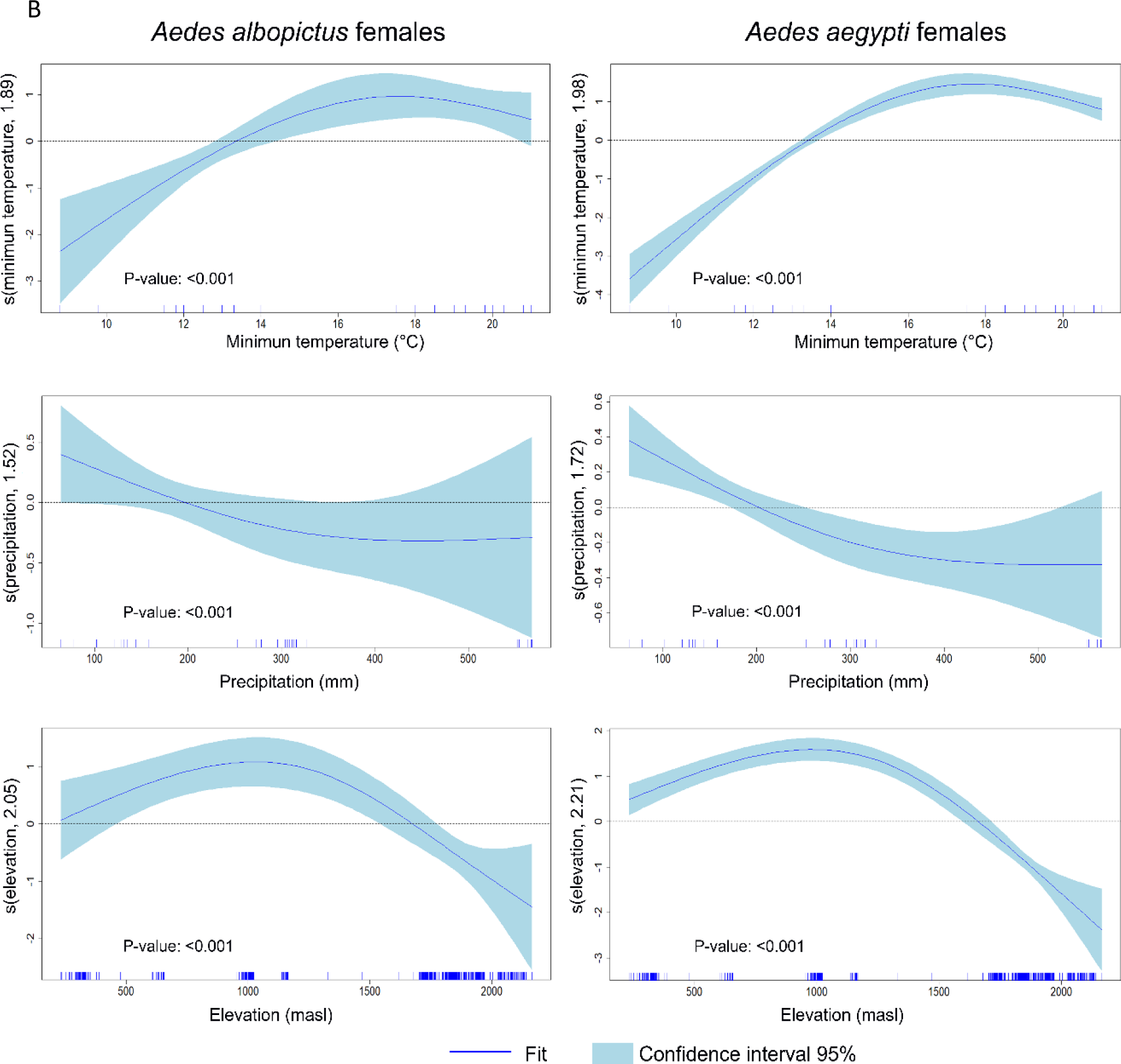
Graphical representation of the estimated results of the GAM model analyzing the climatic factors regarding the number of larvae and females recollected per household. Each panel of the GAM plot displays a smooth function of one predictor, illustrating how variations in that predictor influence the response variable (larvae or females number collected per each *Aedes* species), while the y-axis indicates the predictor’s effect, interpreted as deviations from the average of the response variable, with zero indicating no effect and deviations indicating positive or negative influences. Each curve represents a non-parametric estimate of the predictor effect on the response, with shaded blue light areas indicating the 95% confidence interval, where a wider interval signifies more uncertainty. Additionally, the rug plot on the x-axis shows the observed data points’ distribution, with clusters of tick marks highlighting areas of higher data concentration. A panel indicates the number of collected larvae per house as the response variable. B panel shows the number of females collected on each house, for both *A. albopictus* and *A. aegypti*.

Larval numbers of *A. aegypti* were positively associated with households in the highest and lowest levels of smoothed terms for house conditions and those with little to no knowledge about dengue and its vector (Fig 4). Conversely, larval numbers of *A. albopictus* increased in households with poor house conditions (those constructed mainly with wood and no public service availability) (S1 Fig S1) and those with medium to low knowledge of dengue and the vector (Fig 4).

**Figure 4.**
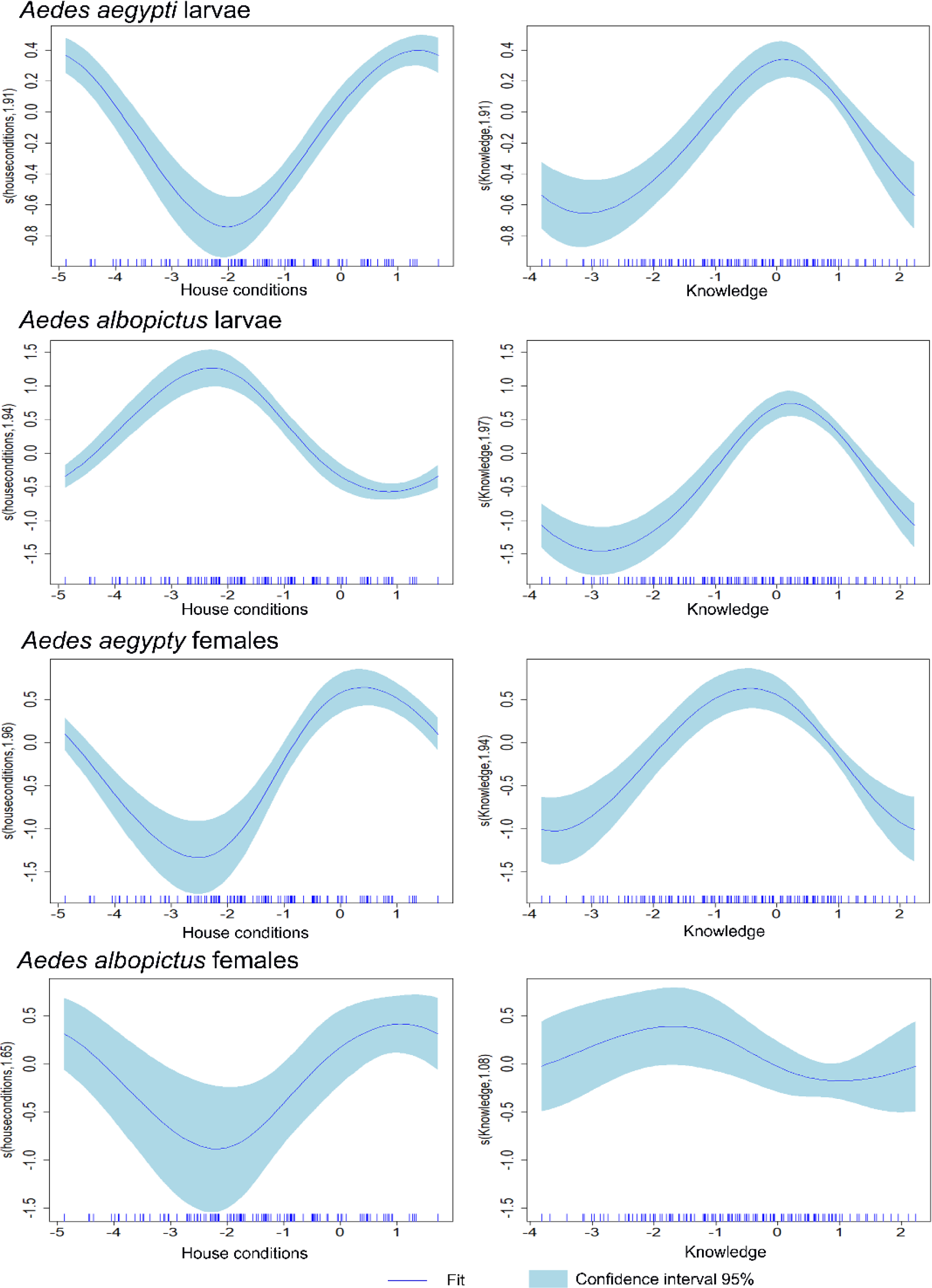
Graphical representation of the estimated results of the GAM model analysis for the reduced anthropogenic variables termed as house conditions and knowledge, against the number of larvae and females recollected at each household. Each panel of the GAM plot displays a smooth function of one predictor, illustrating how variations in that predictor influence the response variable (larvae or females number collected per each *Aedes* species), while the y-axis indicates the predictor’s effect, interpreted as deviations from the average of the response variable, with zero indicating no effect and deviations indicating positive or negative influences. Each curve represents a non-parametric estimate of the predictor effect on the response, with shaded blue light areas indicating the 95% confidence interval, where a wider interval signifies more uncertainty. Additionally, the rug plot on the x-axis shows the observed data points’ distribution, with clusters of tick marks highlighting areas of higher data concentration.

### Adults

#### Adult mosquitoes

In total, 500 adult *A. aegypti* (110 from rural and 390 from urban areas) and 104 adult *A. albopictus* (49 from rural and 55 from urban areas) were collected. Of these, 238 and 86 were female *A. aegypti* and *A. albopictus*, respectively. Eight female and one male adult *A. albopictus* were found exclusively in the extra-domicile environments in the rural areas of Patía and Popayán. Generalized linear models showed that the presence of female *A. aegypti* was positively associated with urban areas, whereas the abundance of female *A. albopictus* was linked to the number of water containers per household (Table 2).

Both species exhibited similar patterns for climatic variables. The number of adult female mosquitoes increased with the smoothed term for minimal temperatures above 15 °C, peaking approximately at 18–19 °C, and then declining at 21 °C. Precipitation > 200 mm negatively affected female abundance. Elevations between 100–1700 masl were associated with a relatively high number of females collected for both the *Aedes* species (Fig 3B). Household-level factors also influenced the abundance of adult females. High house conditions were positively associated with the number of female mosquitoes for both species; however, for *A. albopictus*, the number of females increased even under the lowest house conditions. Female *A. aegypti* was highly abundant in households where inhabitants lacked knowledge about dengue and its vectors, whereas female *A. albopictus* was more abundant in households with accurate knowledge about dengue and its vector (Fig 4).

### Risk of mosquito biting

Changes were observed in the risk of mosquito biting among the study municipalities: in the case of *A. aegypti*, a high risk for mosquito biting, measured by the density of females recollected per house inhabitant, was observed in the rural and urban areas of Patía; however, relatively high pupae per person index was observed in the urban areas of Piamonte (Fig 5). For *A. albopictus*, the risk of biting by both females and pupae per person was lower than that for *A. aegypti*, except for the rural areas in Popayán (Fig 5).

**Fig 5.**
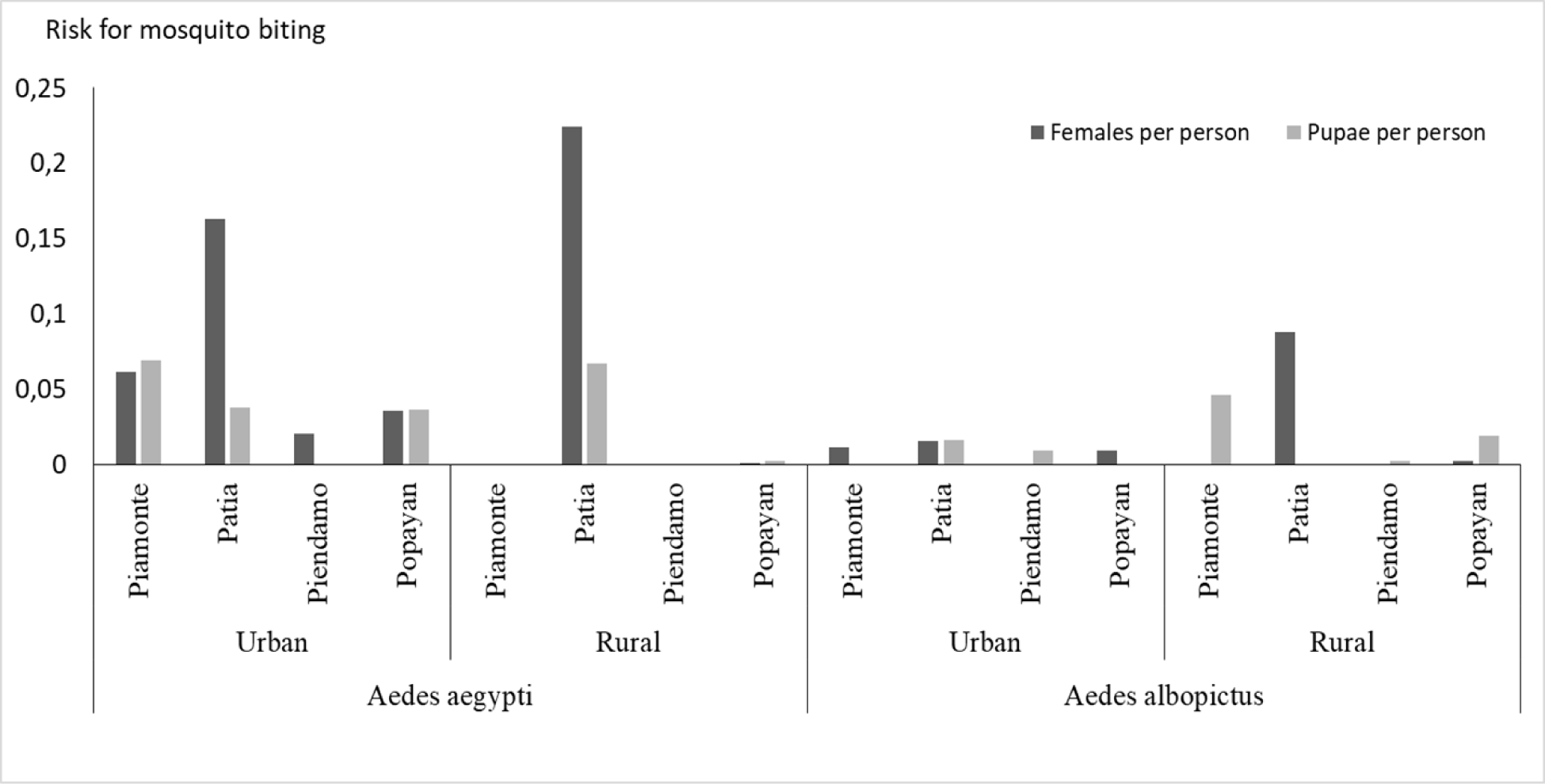
Estimated rate for mosquito biting at each visited house, measure by the pupae per person index (number of pupae collected/number of house inhabitants) and by female per person (number of collected females/number of house inhabitants). Rates were measured for the urban and rural areas of each municipality for both *A. aegypti* and *A. albopictus* species.

### Arbovirus infection

We detected arbovirus infections in 23% (21/89) *A. aegypti* pools with a minimum infection rate (MIR) of 94.38), of which 12.4% (11/89) were positive for DENV (MIR, 41.2) and 12.4% (11/89) for CHIKV (MIR, 49.4). For *A*. *albopictus,* as their abdomens or thorax were individually processed for the female mosquitoes, the infection rate as the percentage of infected abdomens was 61.7% (21/34), with 41.2% (14/34) by CHIKV and 23% (8/34) by DENV; no ZIKV infection was detected in females; however, ZIKV was found in male mosquitoes, indicating virus circulation (Table 3). For *A. aegypti*, the highest infection rate was found in the urban area of Piamonte (MIR, 171.4), followed by that of the urban areas of Patía (MIR, 115.8) and Popayán (MIR, 133.3). The most common natural infection in Patía was by CHIKV in both urban and rural areas, whereas DENV infection was predominant in Piamonte and Popayán. In female *A. albopictus* from Patía, Piendamó, and Popayán, CHIKV was the most frequent infection, except in urban Popayán where only one female mosquito was positive for DENV (Table 3).

**Table 3.**
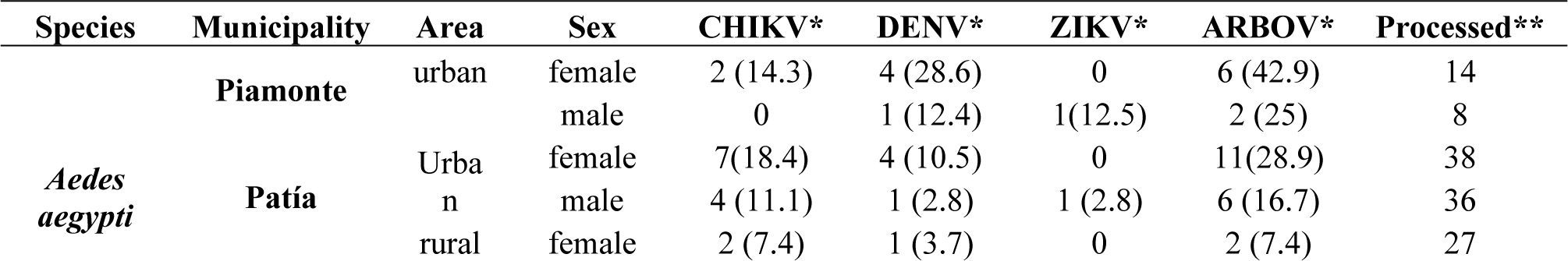
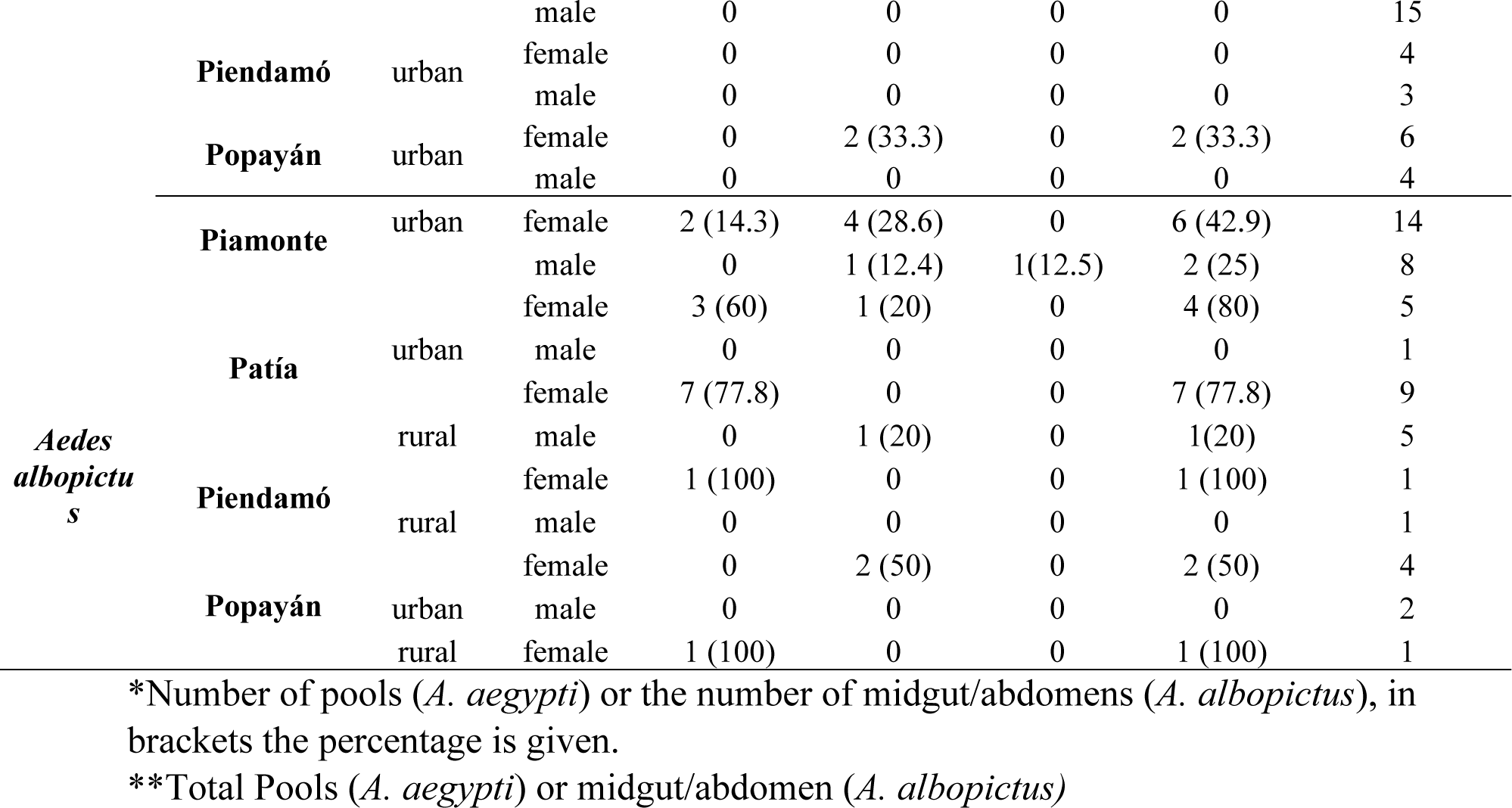
DENV, ZIKV or CHIKV infection in *A. aegypti* pools and *A. albopictus* midguts or abdomens.

We found evidence of DENV and CHIKV dissemination in mosquito tissues, as observed by RNA amplification in the legs or thorax of female *A. albopictus* (Table 4), indicating that the arboviruses escaped the midgut barrier. Furthermore, evidence for the infection by DENV (serotype 2) and CHIKV was indicated by positive amplification in the salivary glands, suggesting the putative role of *A. albopictus* as a vector for both DENV and CHIKV.

**Table 4.**
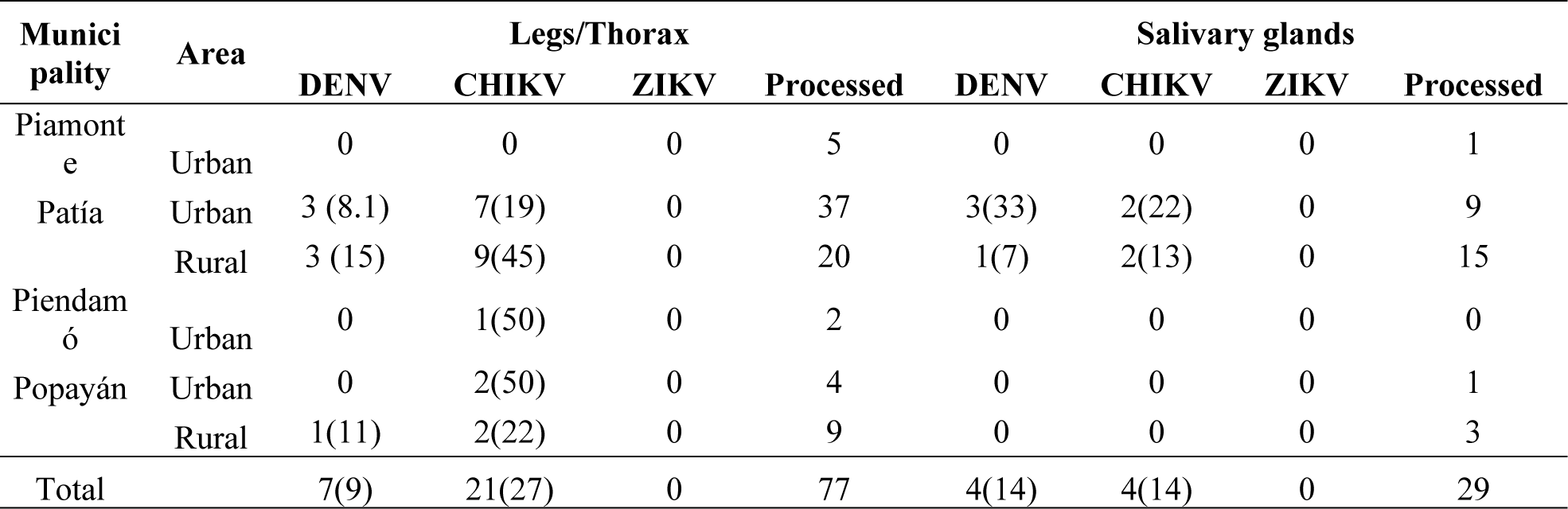
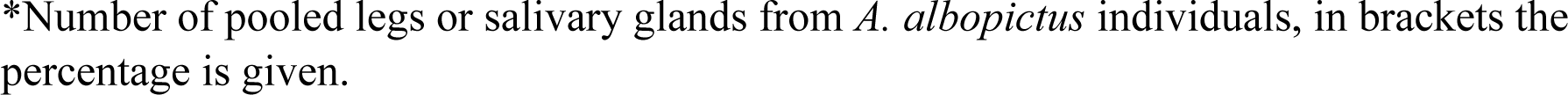
Arbovirus dissemination.

| Municipality | Area | Legs/Thorax |  |  |  | Salivary glands |  |  |  |
| --- | --- | --- | --- | --- | --- | --- | --- | --- | --- |
|  |  | DENV | CHIKV | ZIKV | Processed | DENV | CHIKV | ZIKV | Processed |
| Piamonte | Urban | 0 | 0 | 0 | 5 | 0 | 0 | 0 | 1 |
| Patía | Urban | 3 (8.1) | 7(19) | 0 | 37 | 3(33) | 2(22) | 0 | 9 |
|  | Rural | 3 (15) | 9(45) | 0 | 20 | 1(7) | 2(13) | 0 | 15 |
| Piendamó | Urban | 0 | 1(50) | 0 | 2 | 0 | 0 | 0 | 0 |
| Popayán | Urban | 0 | 2(50) | 0 | 4 | 0 | 0 | 0 | 1 |
|  | Rural | 1(11) | 2(22) | 0 | 9 | 0 | 0 | 0 | 3 |
| Total |  | 7(9) | 21(27) | 0 | 77 | 4(14) | 4(14) | 0 | 29 |

\*Number of pooled legs or salivary glands from *A. albopictus* individuals, in brackets the percentage is given.

## DISCUSSION

Our detailed and integrative analysis of ecological, environmental, and sociodemographic factors, combined with molecular detection and mosquito dissections, helped us establish detailed information about the ecoepidemiology of Aedes-borne viruses in the study area. *A. aegypti* and *A. albopictus* were able to colonize areas up to 2100 masl in Colombia utilizing artificial containers as shared breeding sites in urban and rural habitats. The presence of these species was differentially explained by combining some climatic and anthropogenic factors, with the latter being analyzed and integrated using principal component analysis to obtain informative variables related to household conditions and knowledge, it is worth mentioning that a larger sample size is needed to gather more data on the effects of seasonality and climatic variability on the presence of these species. Additionally, we detected DENV, ZIKV, and CHIKV circulating in mosquito vectors, and evidence of viral dissemination and salivary gland infection was found in *A. albopictus* females, indicating its potential role as an arbovirus vector in the study area. However, exhaustive studies under laboratory conditions, focusing on their competence and vectorial capacity, are necessary to gain a clear understanding of the role of these populations as vectors in the study area.

*A. albopictus* commonly inhabits peri- and extra-domicile environments [38]. However, we found *A. albopictus* indoor also, indicating contact with humans. This species showed a strong capacity to colonize rural and urban habitats, as observed elsewhere [22,23]. Natural larval habitats were not as important for *A. albopictus* compared to that observed in other studies reporting that they mostly use natural breeding sites [14]. In our study, both *A. aegypti* and *A. albopictus* were found up to 2100 masl, although *A. aegypti* has been documented at up to 2200 masl in Colombia [39], which is the highest reported elevation for *A. albopictus* in the country. This demonstrates the ability of mosquitoes to colonize areas above 1800 masl as well as urban and rural habitats. However, the larval and adult numbers of both species declined above 1500– 1700 masl, probably because of decreases in minimum temperatures. Similarly, the abundance and activity of *A. albopictus* decrease with increasing elevation in Italy [40]. However, analyzing this together with anthropic conditions, the decrease could be associated with an improvement in house conditions, as Popayán (the highest elevation site) is the major city in the department, with most houses having considerable availability of public services.

Previous studies using GAM models have reported a positive association between female *A. aegypti* infestation and minimum temperature, with collections peaking approximately at 18 °C and declining above 75% relative humidity (da Cruz Ferreira et al., 2017), which is similar to our results. The presence of *A. albopictus* can be considerably predicted by combining climatic variables and neighborhood conditions [41], and we also found that the combination of climatic and anthropic factors provided a considerable understanding of the ecology of both species.

Environmental capital related to house conditions improves the models for presence of *Aedes* mosquito, with households with medium environmental capital having relatively high immature collections [33]; however, this study did not distinguish between *Aedes* species. We found that the larval abundance of *A. albopictus* and *A. aegypt*i increased with the highest and lowest house conditions, respectively; however, this is related to sociodemographic and behavioral conditions [27]; house constructions materials are associated with the presence of mosquitoes indoor, as construction materials other than concrete can relatively easily allow the entrance of mosquitoes growing outside [42]. We also noticed that breeding places for *A. albopictus* were mainly found outside the house.

*A. albopictus* and *A. aegypti* show overlapping distributions and breeding sites, despite of the reports of competition and displacement [43]. In the present study, these species shared some breeding sites without the evidence for competing; however, *A. aegypti* was notably relatively abundant. The differential effects of climate and house conditions on their abundance likely allowed *A. aegypti* and *A. albopictus* to avoid temporal overlap in the study area by partitioning resources. This niche separation may prevent direct competition in terms of populations despite their cooccurrence and shared breeding sites.

Although laundry sinks, the main breeding places for both species, are not directly influenced by rain, small breeding places, specifically important for *A. albopictus*, are influenced by rainy season. The relationship between the presence of mosquitos and rainfall is usually contradictory, since rainfall favors the presence of small mosquito habitats, together with certain house and knowledge conditions, thereby favoring the presence of mosquitoes in artificial water containers at the domicile [44,45]. However, excess rain may flush the water containers, and immature mosquitoes may die [46], which could explain the negative relationship between precipitation and *A. aegypti* larvae. For *A. albopictus*, we found an association with the highest rainfall periods, which could be explained by egg-laying behavior and the ability of this species to hatch during the first immersions [47], which may help explain the cooccurrence of both species in small containers. Because the eggs of *A. aegypti* require different immersions for hatching, creating an asynchrony in using small breeding sites before rain flushes them, this could favored the contribution of *A. albopictus* as an arbovirus vector during inter-endemic periods, sustaining transmission for the longest periods [48].

Regarding natural infections, the rate of DENV infection observed in *A. aegypti* was similar to that previously reported for some endemic in Colombia [49]. Other studies screening pools of female *A. aegypti* for DENV, ZIKV, and CHIKV have reported positive percentages ranging between 0–40.7% for DENV and 0–12.5% for CHIKV [43–46]. Regarding DENV and CHIKV, we found 12.4% of *A. aegypti* females pools infected by DENV and CHIKV, while 41.2% of *A. albopictus* females were infected by CHIKV, which could highlight the efficient vector competence reported for certain *A. albopictus* populations and CHIKV strains [11]. *A. albopictus* was also found to be associated with considerable vertical and horizontal transmission of DENV [41]. CHIKV viruses also use more than one mosquito species as vectors to facilitate its spread in Brazil [50].

The DENV infection rate in *A. aegypti* is an acceptable predictor of dengue cases [51] and during our sampling period, 73 DENV-positive cases were reported in 2021, with 49 in Piamonte, 15 in Patía, and nine in Popayán. In 2022, 53 DENV cases in Piamonte, 12 in Patía, and four in Popayán have been reported [52], indicating concomitant arbovirus circulation in the study area; however, owing to the weaknesses of the case reporting system, distinguishing between the urban or rural origin of the cases and correlating the cases with entomo-virological surveillance were impossible. However, the presence of arboviruses in mosquitoes above 1800 masl and in rural areas are indications of arbovirus circulation and the potential impact on human health.

Furthermore, in 2024, Popayán is facing the highest outbreak of DENV in its history, with 3,045 cases in the 30^th^ week [52] suggesting the efficiency of *A. aegypti* and *A. albopictus* as vectors for DENV despite their low rate of contact with humans, as observed by the pupae per person and female per person indices [31]; as this is an important predictor for transmission as part of arbovirus Ro [53], however, they could be associated with high presence of DENV in Popayán is their other parameters are high enough.

Only one confirmed case regarding CHIKV and ZIKV was reported in Popayan between 2021– 2022. However, no active detection program for these arboviruses has been reported in Colombia. Additionally, although *A. albopictus* and *A. aegypti* in this study were not tested for the presence of the Mayaro virus (MAYV), Brazilian populations of these mosquitoes have considerable vector competence for MAYV [48], and seroprevalence of MAYV has been reported to be circulating in some municipalities of El Cauca [54]; therefore, these mosquitoes could be participating in the transmission of other arboviruses such as MAYV.

Overall, our results show the integration of environmental, virological, and anthropogenic variables is important for understanding the transmission dynamics and designing effective surveillance and control strategies to prevent the presence of vectors and arbovirus transmission. that the coexistence of two vector species with different ecological characteristics contributes to a complex scenario of arbovirus transmission by expanding its geographic and temporal range, Furthermore, despite *A. albopictus* usually being considered a secondary vector, the results highlight its potential role as a vector in both urban and rural areas, as well as across a wide altitudinal range.

## ACKNOWLEDGEMENTS

We want to thank the administrative staff of Secretaría de Salud del Cauca, specially to Dr. Duban Quintero and Dr. Anderson Piamba. To the Piamonte, Patía, Piendamó, and Popayán mayors’ office, and to their local health secretaries for their logistic support during the fieldwork. We want to thank specially to our fieldworkers Erica Pame, Elvesio Narváez, Victor Perdomo, Cristian Alegría, and Jarvis López. To the local community from urban and rural areas of Piamonte, Patía, Piendamó, and Popayán for receiving us and allowing us to carry out our research. We would like to thank Editage (www.editage.com) for English language editing.

## AUTHOR BIO

Medical entomologist at Instituto de Salud y Ambiente, Vicerrectoría de Investigaciones, Universidad El Bosque, my primary research interests is about vector ecology, vector competence, and vector-pathogen interactions.

## Supporting information captions

**S1 Fig.**
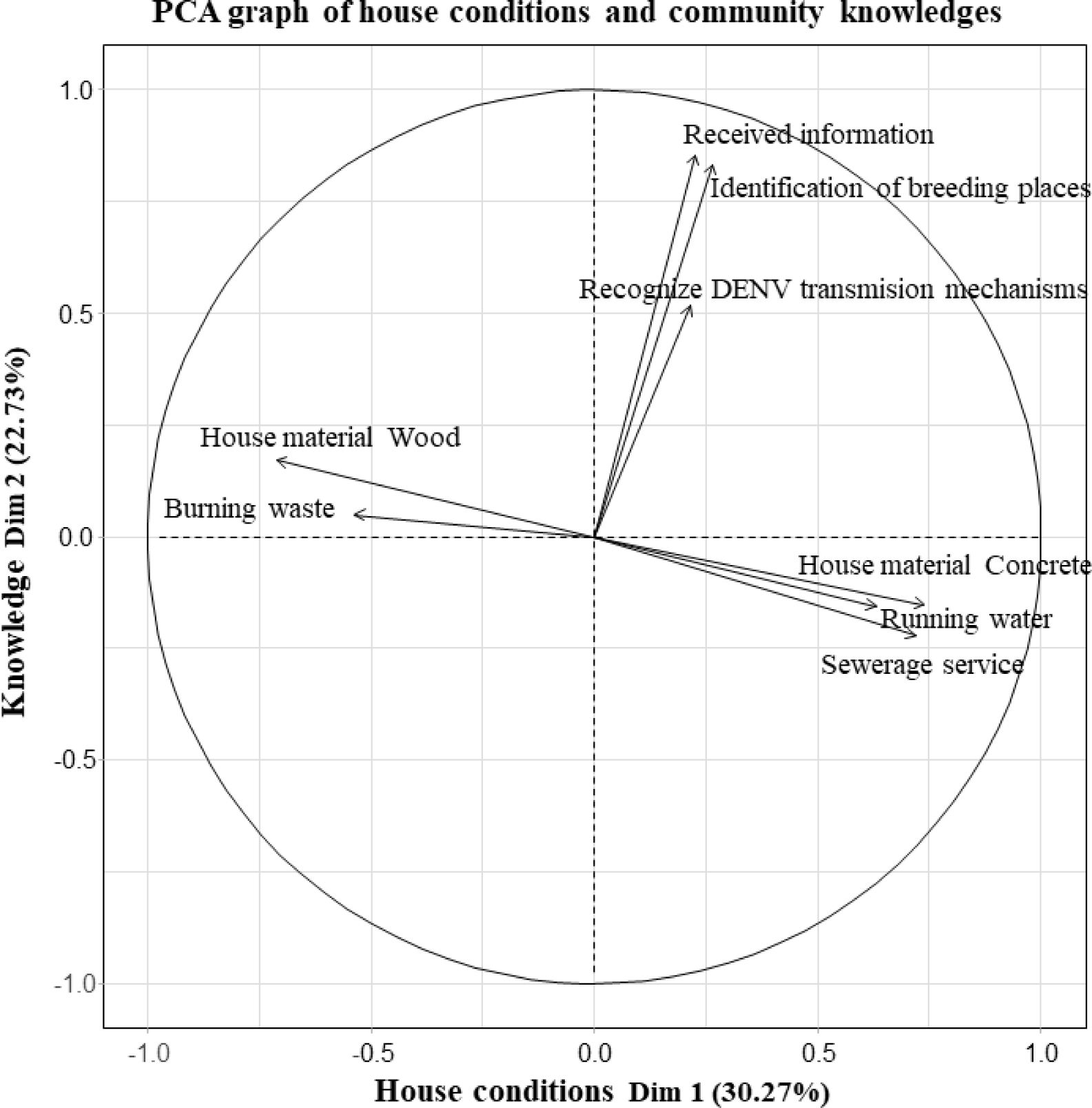
Biplot of principal component analysis. The best-represented variables at each component were selected according to the eigenvalues.

**S1 Table.**
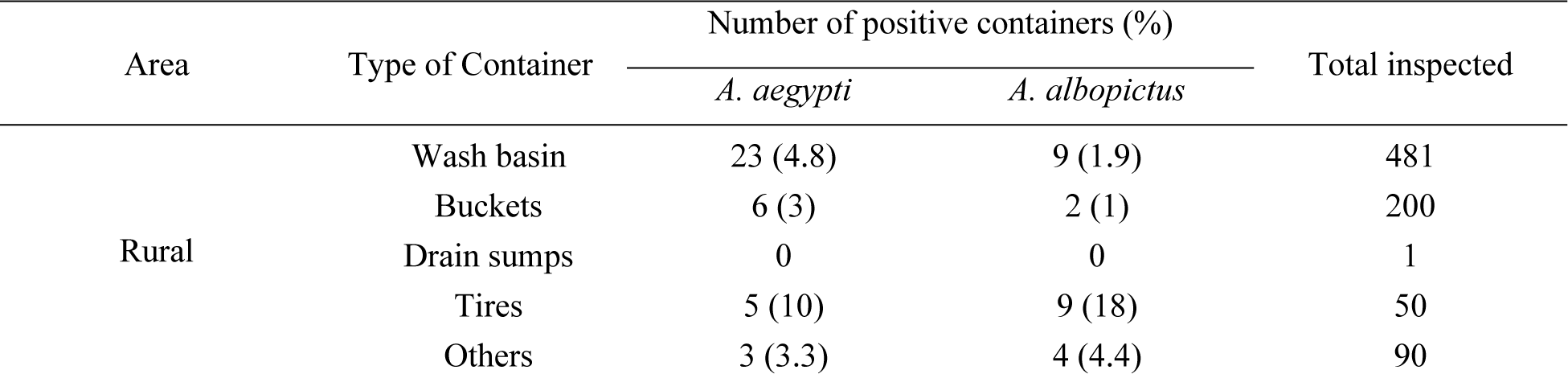
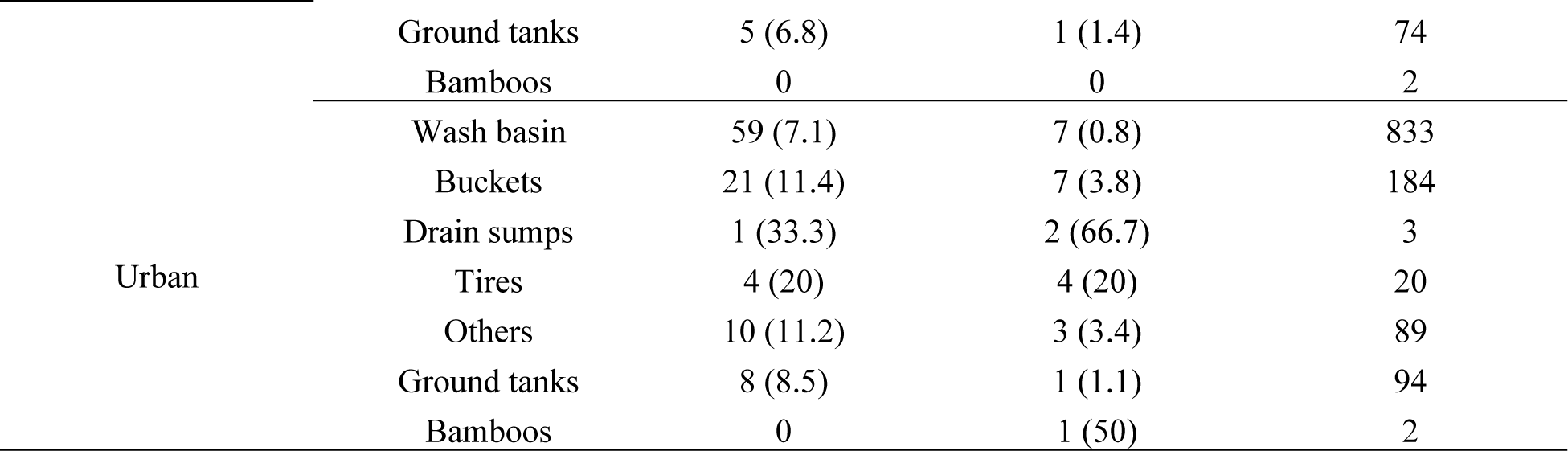
Water containers were found and inspected in urban and rural areas, emphasizing positive breeding places for *A. aegypti* and *A. albopictus*.

